# Evaluation of Conditional Treatment Effect of Salt Stress on Tomato Sugar Content Using Causal Machine Learning: A Pilot Study

**DOI:** 10.1101/2025.07.18.665484

**Authors:** Isao Goto, Shizuka Abiko, Shiori Sugiura, Ai Furudate, Airi Suzuki, Aki Hayashi, Daiki Suzuki, Kaori Kikuchi

## Abstract

Exposing tomatoes to salt stress has been reported to increase the fruit sugar content (°Brix); however, this treatment’s causal impact under varying environmental conditions remains unclear. In this pilot study, a causal inference analysis was conducted using a Causal Tree to analyze the factors that influence the effect of salt-stress treatment on improving the °Brix in tomatoes. Data were collected from a single greenhouse using one cultivar over multiple cultivation periods, totaling 707 fruits. Propensity score matching was applied to reduce the covariate imbalance between the salt (NaCl) treatment and control groups. Using a Causal Tree, conditional average treatment effects were then estimated to assess heterogeneity in treatment impact based on environmental variables, such as temperature, humidity, and illumination. Treatment with NaCl significantly increased °Brix compared to the control, but the magnitude of the effect varied depending on the environmental conditions. The Causal Tree analysis based on the accumulated cultivation data identified specific combinations of environmental factors under which the impact of NaCl treatment on °Brix enhancement was more pronounced. For example, the estimated Conditional Average Treatment Effect (CATE) reached as high as 0.88 when humidity was ≥75%, illuminance was ≥8144 lx, and categorical variable C3 was F1 or F2. In contrast, the CATE dropped to as low as 0.036 under low humidity (<79%) and high temperature (<24°C). These findings highlight humidity as the most influential factor, whose impact was modulated by interactions with temperature and light conditions.

## Introduction

Tomatoes are one of the most important crops in the world, with a global production of over 180 million tons in 2021 [1–2]. They are a valuable source of vitamins, minerals, and antioxidants and are consumed in various forms, including fresh, processed, and canned [3]. Sugar content is considered one of the most important factors affecting tomato fruit quality and consumer satisfaction [4]. Tomatoes with a high sugar content are popular in Japan, China, and Korea, and in Japan in particular, they are considered luxury items and are often traded at high prices. Therefore, a cultivation method that applies water stress to concentrate components within the fruit is used to increase the sugar content in tomatoes [5–6].

In hydroponic cultivation, sodium chloride is added to the nutrient solution to increase the osmotic pressure in the root zone, thereby restricting water absorption [7–8]. Applying salt stress, rather than limiting irrigation, is considered a more efficient cultivation method because it provides uniform water stress across the entire root zone and allows for more precise adjustment of stress levels. However, even if constant salt stress is applied during cultivation, the fruit sugar content is not constant. Even when cultivated without stress, the sugar content of tomatoes fluctuates widely throughout the year [9], largely due to the strong influence of environmental factors on both sugar accumulation and stress responses. Factors that determine the sugar content of fruits include photosynthesis, which synthesizes sugar; respiration, which consumes sugar; and the amount of water supply, which affects sugar concentration. Environmental conditions, such as temperature, solar radiation, and humidity, and the duration of exposure to them, significantly influence these factors. In particular, temperature and solar radiation are important factors that determine the balance between photosynthesis and respiration, and humidity affects water absorption and sugar concentration [10–11]. Moreover, these factors do not act independently but interact, making the relationship complex [12]. Therefore, to stabilize sugar content, it is necessary to analyze in detail how environmental factors such as temperature, solar radiation, and humidity interact with photosynthesis, respiration, and water uptake throughout the seasons. It is also important to understand how these interactions vary across different growth stages and durations, leading to fluctuations in sugar content, to establish management methods based on this analysis.

In this study, tomatoes were cultivated year-round and causal inference was used to examine the treatment effect of salinity stress on sugar content. Specifically, tomatoes were cultivated in nutrient solutions at different times after adjusting for the sowing time. Two groups were established for each sowing period: a salt-stress treatment group and a control group (without salt stress). For tomatoes matured under the same cultivation conditions (environment), the percentage of tomatoes with a sugar content above 6 °Brix (or °Bx, i.e., the percent by weight of sugar solids in a pure sucrose solution) was calculated for both treatment groups. Factors influencing the percentage difference between the two groups were analyzed using causal inference. Based on these results, we aimed to identify environmental factors and tree conditions that enhance the effects of salt stress.

## Materials and methods

### Plant materials and growing conditions

Tomato (*Solanum lycopersicum* L. “Momotaro York”; Takii & Co., Ltd., Japan) seeds were sown nine times between March 2016 and June 2022 (Table 1). The seeds were sown in rockwool plugs (Kiemplug; Grodan Delta, The Netherlands) and allowed to germinate in a temperature-controlled growth chamber at 25/20 ℃ (day/night). After germination, seedlings were transplanted into rockwool cubes (75 × 75 × 65 mm; Grodan Delta) and allowed to grow in a greenhouse supplied with a commercial nutrient solution, Otsuka A (OAT Agrio Co., Ltd., Japan), adjusted to an electrical conductivity (EC) of 0.8 dS/m and pH of 6.0–6.5. Just before first-truss flowering, young plants were transplanted into a nutrient film technique system supplied with the commercial nutrient solution Otsuka SA, adjusted to an EC of 1.2 dS/m. Once the first flower opened on the first truss of each plant, fruiting was facilitated by applying 2-methyl-4-chlorophenoxyacetic acid. The number of flower settings was left to nature. All lateral shoots were removed as they appeared, and the plants were pinched above the third truss with two true leaves over the truss. Cultivation was terminated after the fruits were harvested from the third flower cluster. Cultivation periods A to I are defined in Table 1, based on the sowing dates. The dates when flowers of the first, second, and third trusses began anthesis and the cultivation termination dates are shown for each cultivation period. The dates of anthesis for each truss represent the average flowering dates of the 12 plants surveyed in each cultivation period. During all cultivation periods, ventilation was performed when the temperature exceeded 25 °C in summer, and heating was applied to maintain a minimum temperature of 10 °C in winter. The temperature, relative humidity, and illuminance were measured every 10 min using a data logger (RS-13L; ESPSC MIC Co., Ltd., Japan). Temperature and humidity sensors were installed in a ventilation pipe placed inside the canopy, and the illuminance sensor was installed at the top of the cultivation shelf (height 1.8 m). The average temperature and relative humidity were calculated for each fruit over the period from anthesis to harvest. The illuminance measurements taken every 10 min were converted to a per-second basis, and the integral illuminance was calculated for the period from anthesis to harvest. The environmental data (temperature, humidity, and illuminance) for each fruit were used as covariates (X) in the causal inference analysis, as was the period from anthesis to harvest for each fruit, as shown in Table 2.

**Table 1.**
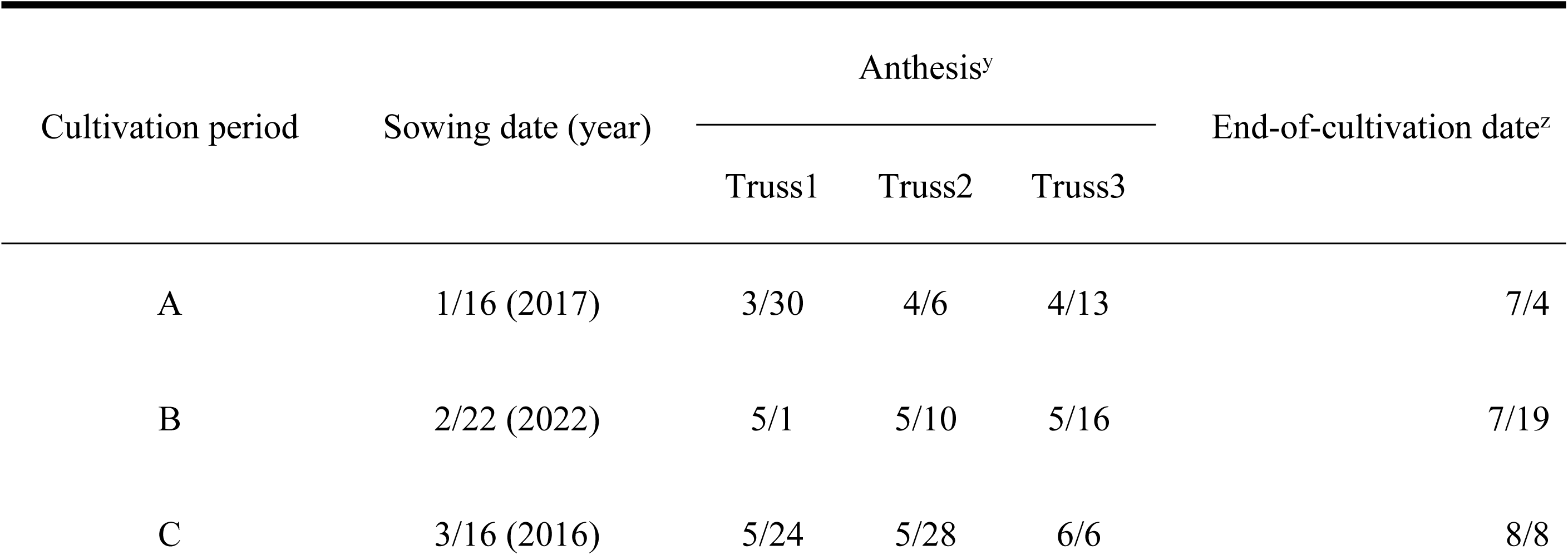
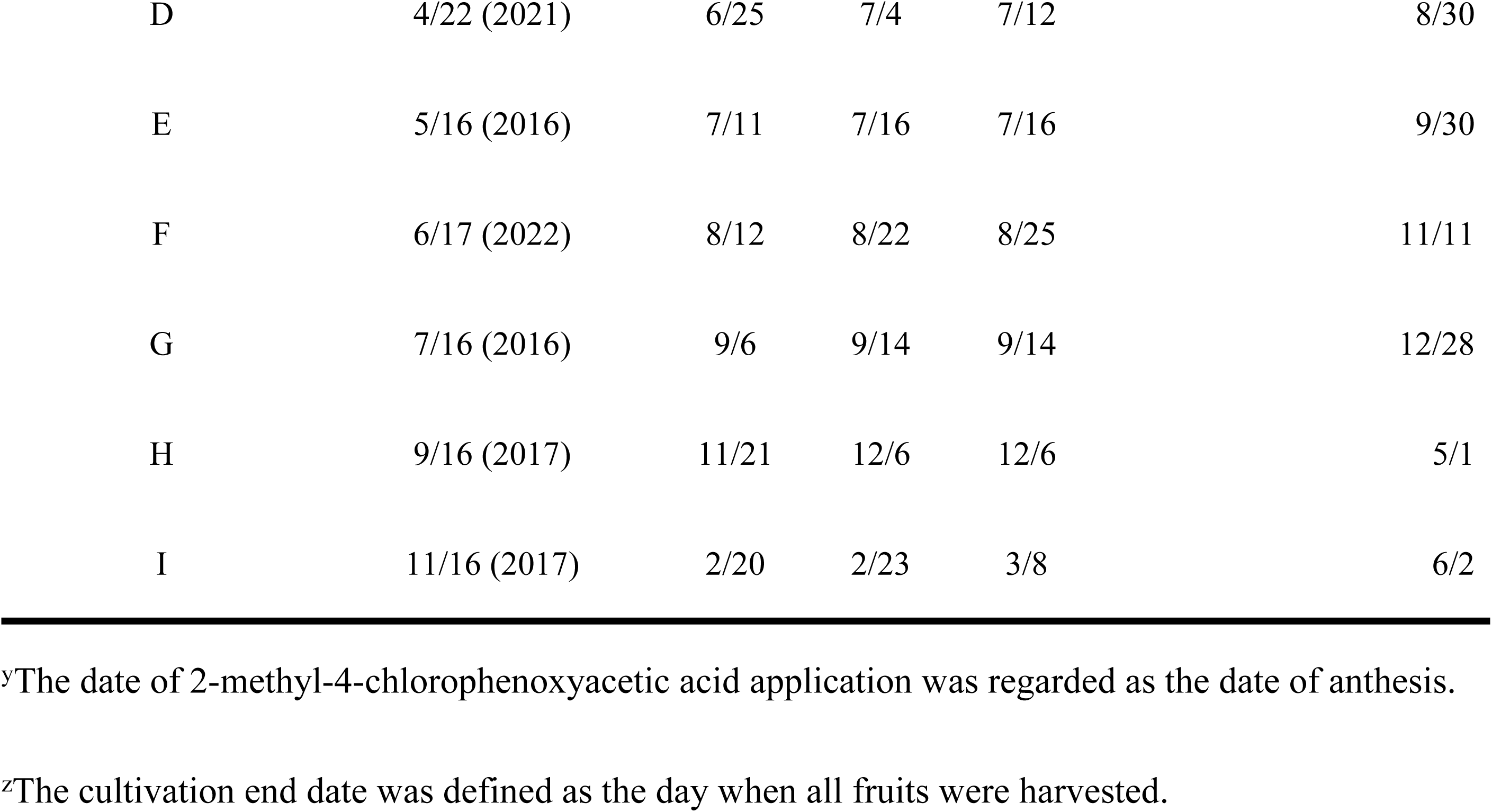
Sowing dates and average anthesis and cultivation end dates (n = 12) for each cultivation period.

**Table 2.**
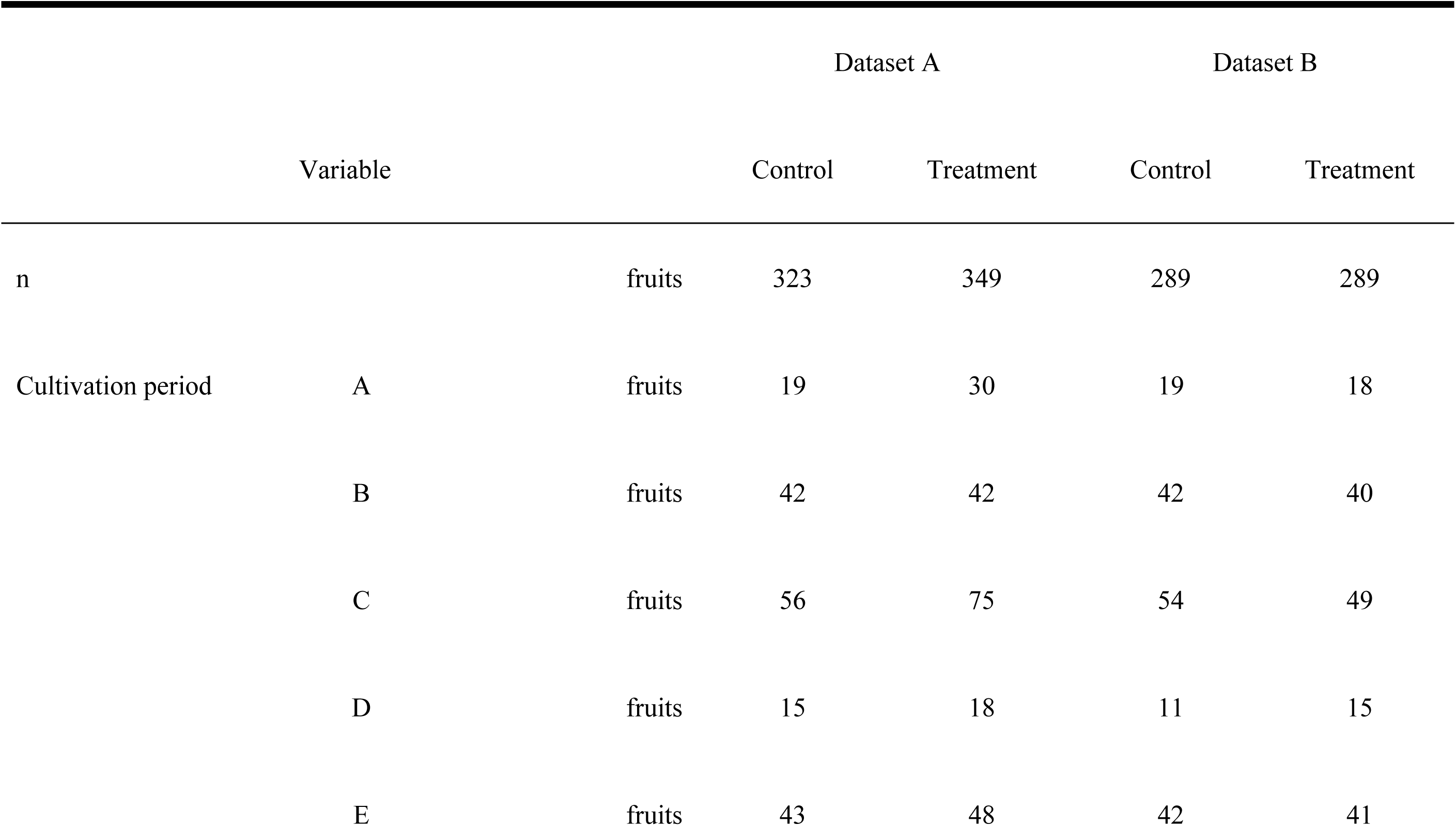
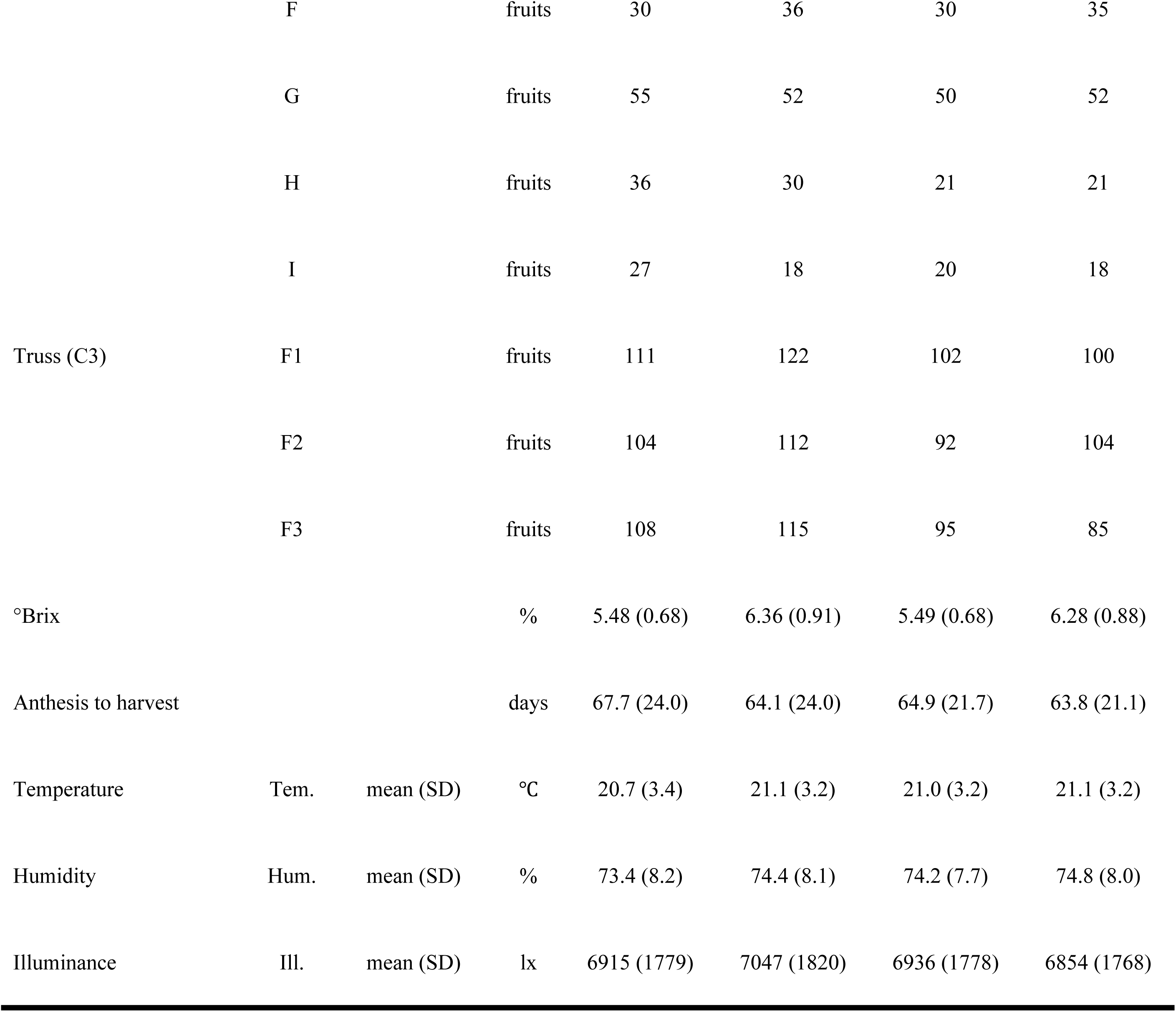
Dataset characteristics before and after propensity score matching.

|  |  |  | Dataset A |  | Dataset B |  |
| --- | --- | --- | --- | --- | --- | --- |
|  | Variable |  | Control | Treatment | Control | Treatment |
| n |  | fruits | 323 | 349 | 289 | 289 |
| Cultivation period | A | fruits | 19 | 30 | 19 | 18 |
|  | B | fruits | 42 | 42 | 42 | 40 |
|  | C | fruits | 56 | 75 | 54 | 49 |
|  | D | fruits | 15 | 18 | 11 | 15 |
|  | E | fruits | 43 | 48 | 42 | 41 |

|  |  |  |  |  |  |  |  |
| --- | --- | --- | --- | --- | --- | --- | --- |
|  | F |  | fruits | 30 | 36 | 30 | 35 |
|  | G |  | fruits | 55 | 52 | 50 | 52 |
|  | H |  | fruits | 36 | 30 | 21 | 21 |
|  | I |  | fruits | 27 | 18 | 20 | 18 |
| Truss (C3) | F1 |  | fruits | 111 | 122 | 102 | 100 |
|  | F2 |  | fruits | 104 | 112 | 92 | 104 |
|  | F3 |  | fruits | 108 | 115 | 95 | 85 |
| °Brix |  |  | % | 5.48 (0.68) | 6.36 (0.91) | 5.49 (0.68) | 6.28 (0.88) |
| Anthesis to harvest |  |  | days | 67.7 (24.0) | 64.1 (24.0) | 64.9 (21.7) | 63.8 (21.1) |
| Temperature | Tem. | mean (SD) | °C | 20.7 (3.4) | 21.1 (3.2) | 21.0 (3.2) | 21.1 (3.2) |
| Humidity | Hum. | mean (SD) | % | 73.4 (8.2) | 74.4 (8.1) | 74.2 (7.7) | 74.8 (8.0) |
| Illuminance | Ill. | mean (SD) | lx | 6915 (1779) | 7047 (1820) | 6936 (1778) | 6854 (1768) |

### Salinity treatments

In the control treatment, the EC level was maintained at 1.2 dS/m from transplantation until the end of cultivation. For the NaCl treatment, NaCl was added to the standard nutrient solution at first-truss flowering and gradually increased to the desired EC of 4.0 dS/m (25 mM NaCl) at second-truss flowering. The EC was maintained at 4.0 dS/m until all the fruits from the third truss were harvested. Previous studies have evaluated the effect of salt concentration on tomato sugar content, with reports indicating that higher salt stress leads to increased sugar content [14–16]. Consequently, many studies have used a salt concentration of 50 mM NaCl (≈ 8 dS/m) to reliably increase the sugar content [17–19]. However, growth and yield significantly decrease when the salt concentration exceeds a certain level [20]. Various strategies have been employed to address this trade-off, such as limiting the timing of salt stress or gradually increasing the salt stress [21].

We aimed not to evaluate the effect of the salt concentration but to elucidate how the cultivation environment influences the effect of salt stress on increasing sugar content. The stress associated with high salt concentrations may be less affected by cultivation conditions; therefore, in this study, we used a lower salt concentration (EC: 4 dS/m, 25 mM NaCl) than typically used in previous studies. We expect the impact of environmental conditions to be more pronounced at this level of salt stress.

### Fruit yield and quality analyses

Marketable fruits with fresh weights of 50 g or more from the surveyed plants’ first, second, and third trusses were selected and harvested when they reached the red stage. After harvesting, the fruits were weighed to determine their fresh weights. The fruits were then split in half. One half was used to measure total soluble solids (°Brix) content using a hand refractometer (PAL-BX|ACOD3; Atago Co., Ltd., Tokyo, Japan) and the other half was oven-dried at 80 °C for 48 h and the dry weight was measured. The total number of sampled fruits was 707, and the Brix values were used for subsequent analyses.

### Analysis

#### Data selection and analysis flow

Dataset A was created by applying basic data cleaning procedures, such as removing missing values, to all fruit (n = 707) and environmental data collected (Fig 1). We examined the data characteristics of Dataset A, including the descriptive statistics of each variable. Propensity score matching was performed to adjust for potential biases between the control and treatment groups, resulting in Dataset B (n = 578). Using Dataset B, the conditional average treatment effects (CATEs) on salt stress were estimated using a Causal Tree.

**Fig 1.**
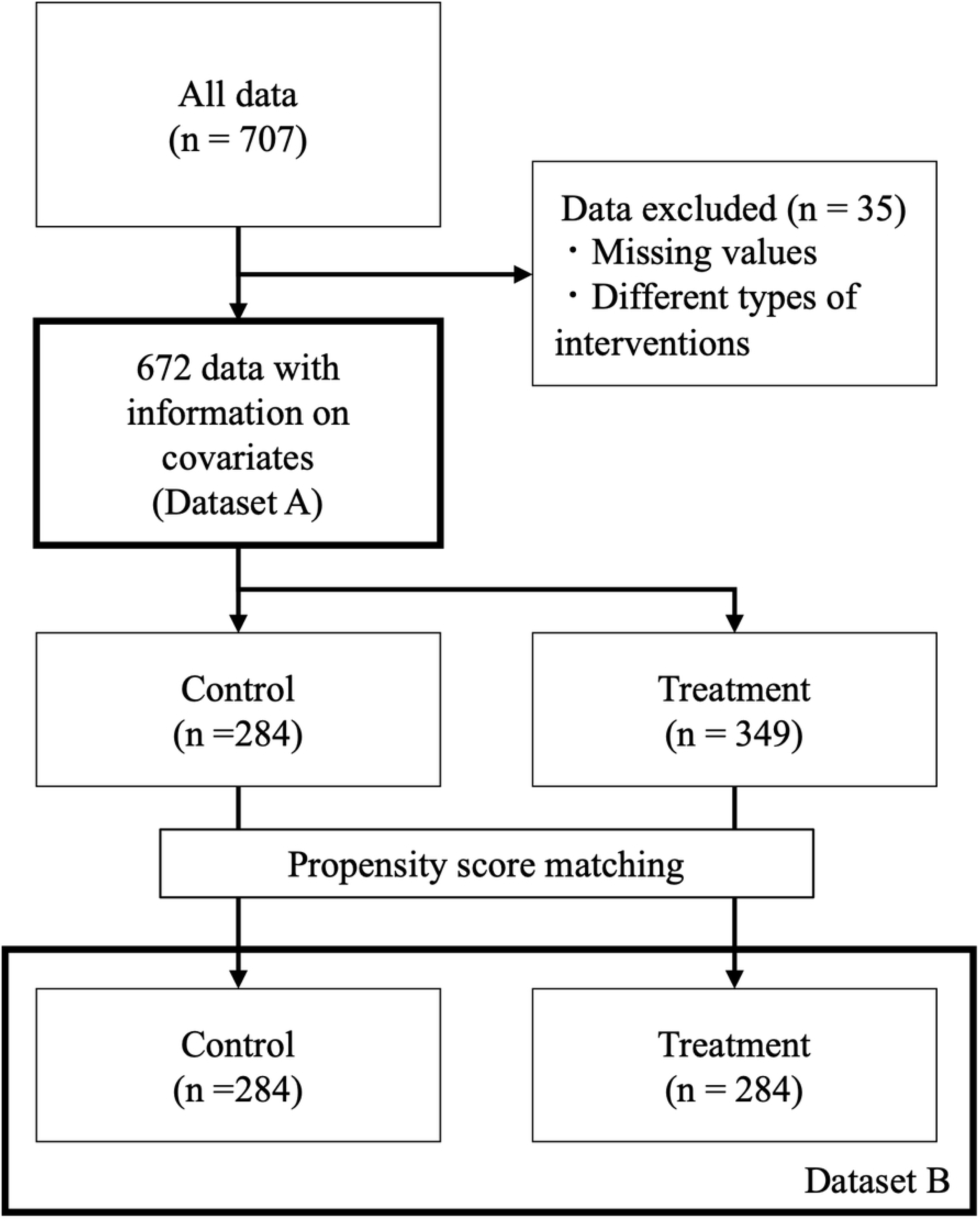
Flow of study data selection. The 672 fruits after data cleaning were prepared as Dataset A and the 578 fruits after propensity score matching as Dataset B.

#### Propensity score matching

We performed 1-to-1 propensity score matching without replacement to pair fruits in the control and treatment groups while adjusting for the covariates listed in Table 2. Propensity scores for treatment were estimated using a logistic regression model. Matching was performed using a caliper width of standard deviations of the logit of the propensity score. A successful balance between the groups was confirmed by an absolute standardized mean difference of < 0.1.

#### Causal Tree

We applied a Causal Tree in this study to estimate the CATE, allowing for heterogeneous NaCl treatment effects [22]. A Causal Tree is a decision tree-based approach that provides interpretable subgroup-level treatment effects and captures nonlinear treatment heterogeneity.

#### Data and variables

- Outcome variable (*Y*): °Brix (1 = above 6%, 0 = below 6%)

In the absence of a widely accepted threshold for this evaluation metric, a provisional cutoff (6%) was determined based on the observed distribution in the treatment and control groups.

- Treatment variable (*W*): NaCl treatment (1 = treated, 0 = control)
- Covariates (*X*): temperature (Tem.), humidity (Hum.), illumination (Ill.), number of trusses (C3; the first, second, and third trusses were designated F1, F2, and F3, respectively)

#### Model specification

The Causal Tree method was implemented using the causalTree package in R (https://github.com/susanathey/causalTree). The hyperparameters were set as follows:

- Splitting rule: Causal Tree (split.Rule = “CT”)
- Cross-validation method: Causal Tree (cv.option = “CT”)
- Minimum leaf size (subgroup size): 30 (minsize = 30)
- Honest splitting: disabled (split honest = FALSE)

The choice of a minimum subgroup size of 30 was based on prior literature, which ensured that each l subgroup contained enough observations for stable effect estimation [22]. Causal Tree-based splitting and cross-validation were used to optimize treatment effect heterogeneity rather than prediction accuracy. Honest splitting was not performed to preserve the entire dataset for estimation purposes, instead of dividing it into separate training and validation sets.

#### Estimation of CATE

The Causal Tree method partitions the data based on covariates *X* and estimates the CATE within each subgroup node using the following equation:

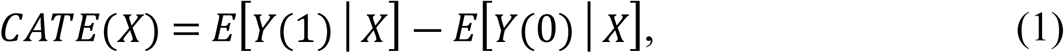

where *E*[*Y*(1)│*X*] represents the expected outcome for the treated group conditioned on *X* and *E*[*Y*(0)|*X*] represents the expected outcome for the control group conditioned on *X*. This approach enables the analysis of how the impact of salt-stress treatment varies across different subgroups, considering factors such as age and environmental conditions.

### Statistical analysis

To compare the means between the two groups, Welch’s t-test was employed because one group did not meet the assumption of normality. The assumption of normality was assessed using the Shapiro– Wilk test. In addition to the p-values, mean differences with 95% confidence intervals were reported to indicate the magnitude and uncertainty of the effect.

### Analytical environment

All analyses were performed using R version 4.2.2 (R Foundation for Statistical Computing, Vienna, Austria). The following R packages were used: effectsize (1.0.0), devtools (2.4.5), tidyverse (1.3.1), Matching (4.10.8), and ggplot2 (3.4.0). We used the causalTree package (version 1.0) developed by Athey and Imbens [22] for model construction. This package was installed in the susanathey/causalTree GitHub repository.

## Results

The sample size for Dataset A was 672, with the control and treatment groups consisting of 323 and 349 samples, respectively (Fig 1). Sample sizes varied across cultivation periods, and the transplant-to-harvest period differed by an average of 2.5 days after adjusting for covariates (Table 2). Propensity score matching was applied to ensure a more balanced comparison, resulting in reduced variation in most covariates between the two Dataset B groups (Table 2). NaCl treatment significantly increased the °Brix compared with the control (p < 0.01). The mean °Brix of the control was 5.48 (95% CI: 5.40– 5.55) and that of salt-stress treatment was 6.38 (95% CI: 6.26–6.45) (Fig 2). The mean difference was 0.88 (95% CI: 0.76–1.00), suggesting a higher sugar content under salt stress.

**Fig 2.**
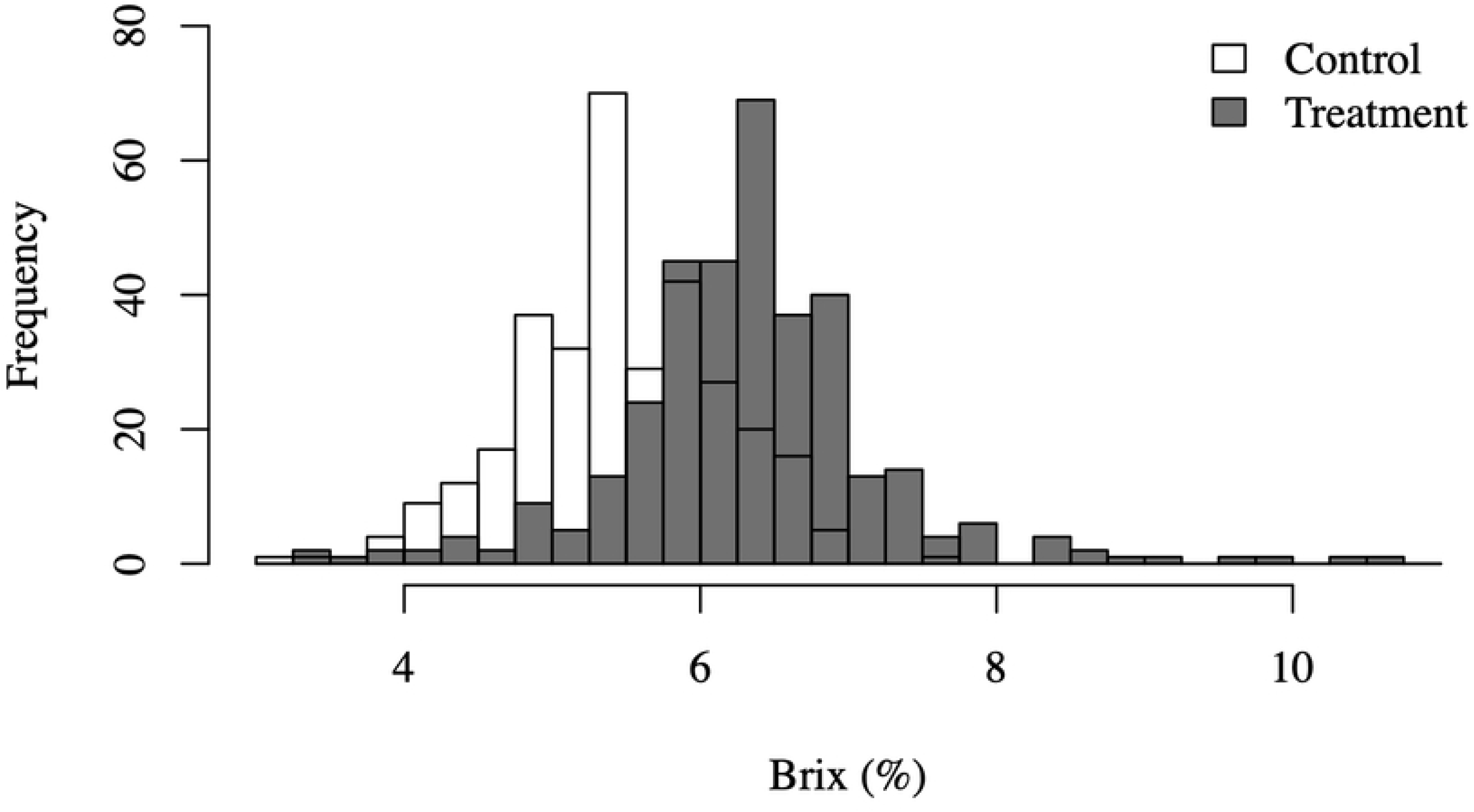
Overlapping histograms showing the distribution of tomato Brix in the control (white) and salt-stressed treatment (gray) groups. The x-axis represents Brix values, and the y-axis indicates the number of fruits (frequency).

We examined whether the effectiveness of the NaCl treatment varied depending on the cultivation period. Figure 3 illustrates the relationship between the salt-stress treatment and °Brix values across different cultivation periods. NaCl treatment consistently increased the fruit °Brix in all cultivation periods; however, the magnitude of its effect varied with the cultivation period. Cultivation periods A, B, C, E, G, H, and I showed meaningful effects, whereas D and F yielded only negligible differences (S1 Table). Table 3 summarizes the mean and standard deviation of the room temperature, humidity, and illumination in the control and treatment groups during each growing period. There were only minor differences in environmental conditions between the control and treatment plots within each period. In contrast, substantial differences in environmental conditions were observed across cultivation periods. For instance, while the average difference in temperature between the control and treatment was less than 0.5 °C in cultivation periods B and C, the difference in humidity exceeded 7%.

**Fig 3.**
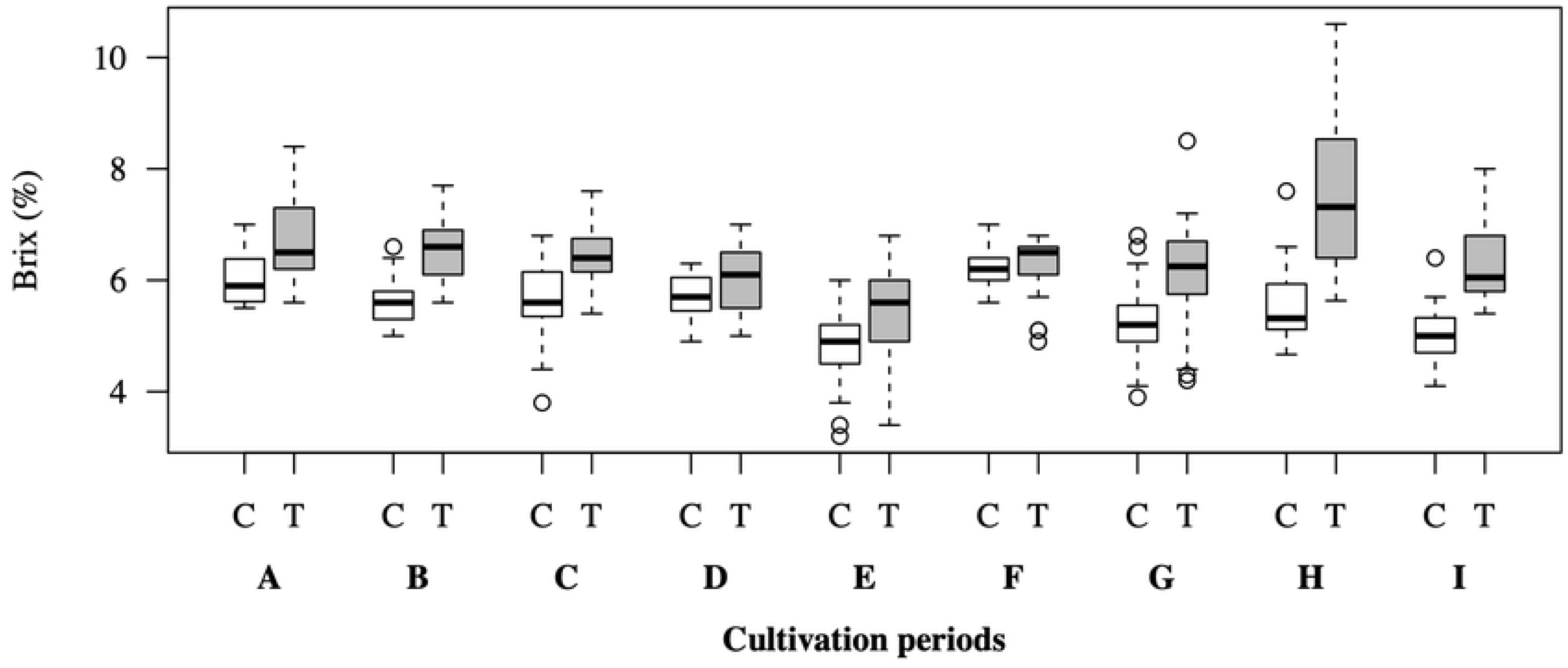
Box plots of tomato °Brix for the control (white) and salt-stressed treatment (gray) groups across nine cultivation periods (A–I). The boxes represent the interquartile range (IQR) from the first (Q1) to the third (Q3) quartile, with the central line indicating the median. Whiskers extend to data points within 1.5 × IQR, and outliers are plotted as circles (○).

**Table 3.**
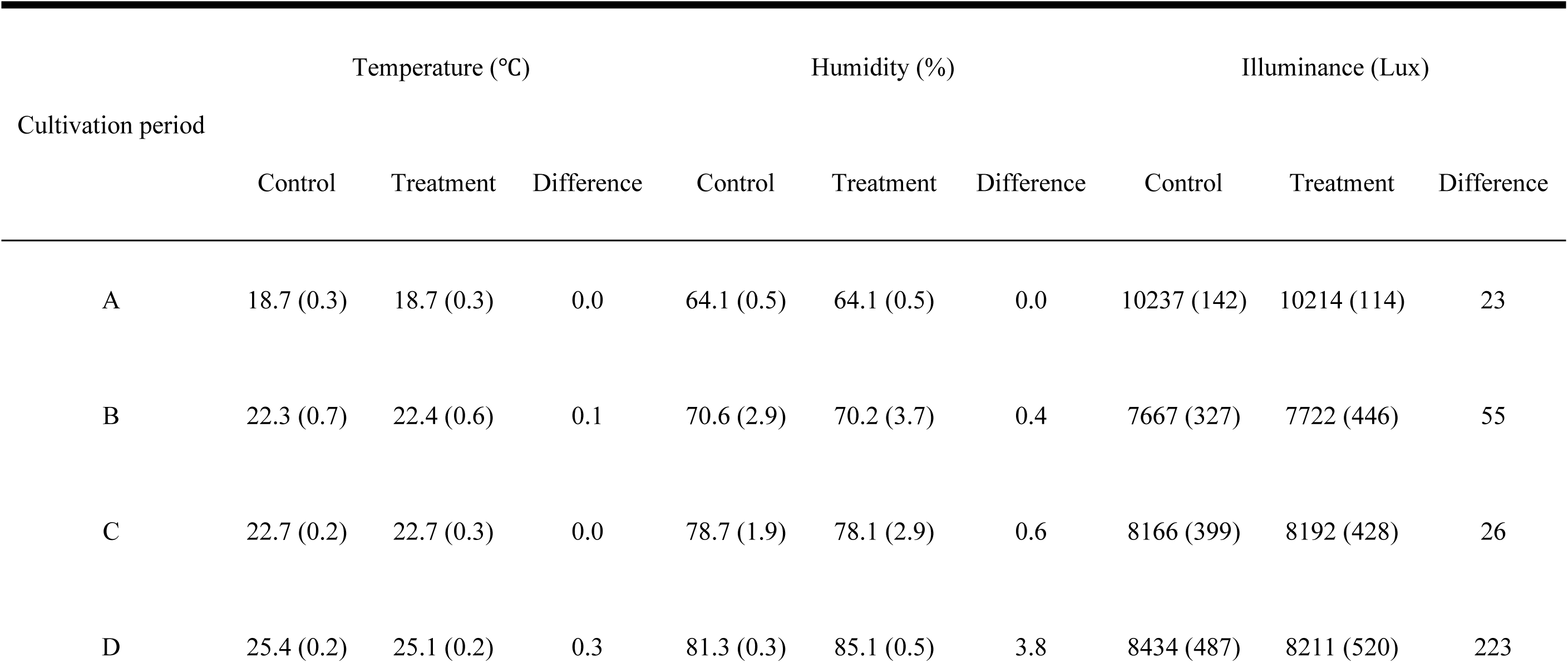
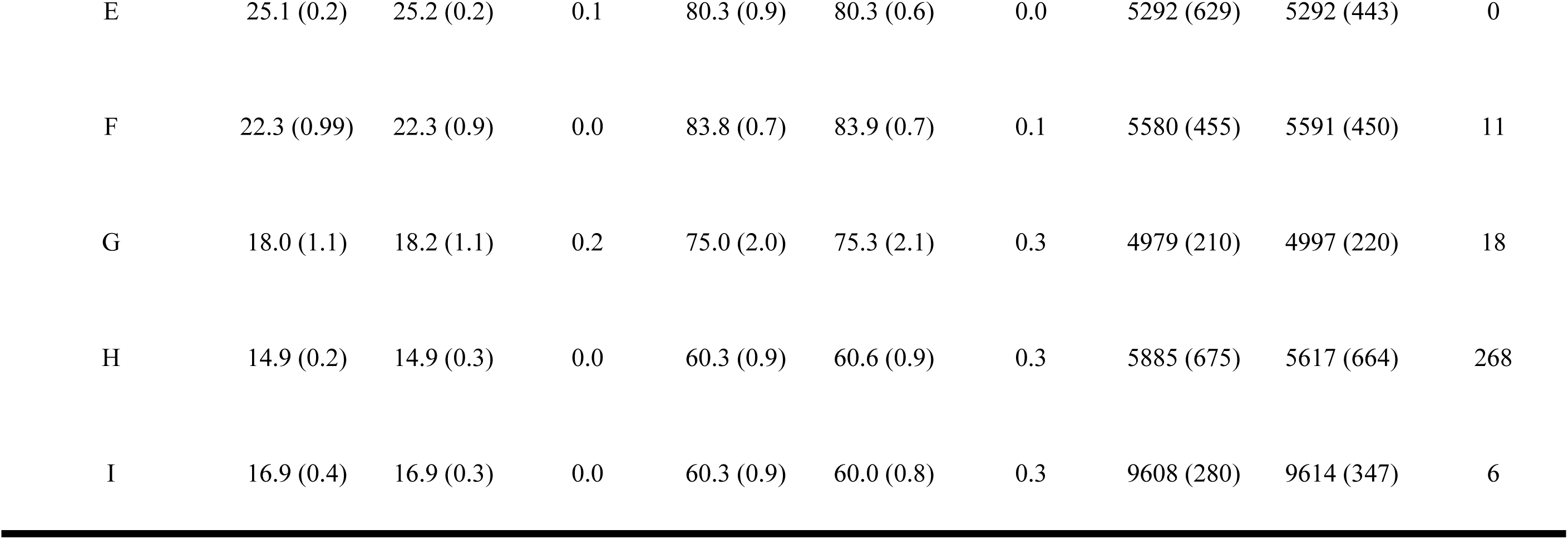
Mean (SD) of temperature, relative humidity, and illuminance in the greenhouse for the control and NaCl treatment groups across each cultivation period and differences between the group means.

**Supplementary table 1.** Difference in average °Brix of tomato fruits between control and treatment.

| Cultivation | Mean | Mean | Mean |  |  |  |  |
| --- | --- | --- | --- | --- | --- | --- | --- |
| period | (Control) | (Treatment) | Difference | p-value | Cohen's d | 95% CI (Lower) | 95% CI (Upper) |
| A | 6.0114 | 6.7600 | 0.7486 | < 0.0001 | 1.2326 | 1.8273 | 0.6270 |
| B | 5.6095 | 6.5762 | 0.9667 | < 0.0001 | 2.2931 | 2.8497 | 1.7275 |
| C | 5.6589 | 6.4600 | 0.8011 | < 0.0001 | 1.4003 | 1.8020 | 0.9930 |
| D | 5.6933 | 6.0167 | 0.3233 | 0.0772 | 0.6288 | 1.3157 | -0.0681 |
| E | 4.8349 | 5.3979 | 0.5630 | 0.0005 | 0.7521 | 1.1733 | 0.3269 |
| F | 6.2267 | 6.2917 | 0.0650 | 0.5295 | 0.1541 | 0.6321 | -0.3251 |
| G | 5.2382 | 6.1750 | 0.9368 | < 0.0001 | 1.3277 | 1.7471 | 0.9030 |
| H | 5.4972 | 7.5856 | 2.0883 | < 0.0001 | 1.9409 | 2.5926 | 1.2746 |
| I | 5.0278 | 6.2972 | 1.2694 | < 0.0001 | 2.0450 | 2.8538 | 1.2145 |

This table summarizes the mean, mean difference, p-value (by Welch's t-test), Cohen's d (standardized mean difference), and 95% confidence interval of the mean difference for each variable for the control and treatment groups at each cultivation period.

We also examined how Causal Tree-derived subgroups corresponded to the cultivation period. While D was assigned to a single terminal node because of its limited sample size, the other periods were represented across multiple subgroups. Furthermore, each Causal Tree subgroup included samples from multiple cultivation periods (Table 4), supporting the interpretation that subgrouping was primarily driven by environmental and contextual variables rather than seasonal classification alone.

**Table 4.** Number of fruits observed during the growing season for each subgroup node in the causal tree.

| Cultivation period | Subgroup |  |  |  |  |  |  |
| --- | --- | --- | --- | --- | --- | --- | --- |
|  | 1 | 2 | 3 | 4 | 5 | 6 | 7 |
| A | 0 | 0 | 8 | 0 | 0 | 29 | 0 |
| B | 0 | 0 | 29 | 0 | 0 | 9 | 44 |
| C | 30 | 0 | 0 | 10 | 62 | 1 | 0 |
| D | 0 | 26 | 0 | 0 | 0 | 0 | 0 |
| E | 0 | 80 | 0 | 0 | 3 | 0 | 0 |
| F | 59 | 6 | 0 | 0 | 0 | 0 | 0 |
| G | 3 | 0 | 30 | 61 | 0 | 0 | 8 |
| H | 0 | 0 | 12 | 0 | 0 | 0 | 30 |
| I | 0 | 0 | 14 | 0 | 0 | 24 | 0 |

The outcome variable was provisionally dichotomized based on °Brix values, classifying fruits as either < 6% or ≥ 6% to evaluate the effect of the salt-stress treatment. First, the CATE was estimated using the Causal Tree method; its structure is illustrated in Fig 4. After propensity score matching, the overall average treatment effect was 0.45. However, the CATE differed across subgroups; for instance, it was 0.13 when the highest splitting variable, humidity, was < 79% and 0.63 when humidity was ≥ 79%. The highest estimated CATE of 0.88 (95% CI: 0.77–0.99) was observed under the following conditions: Hum. ≥ 75%, C3 = F1 or F2, Ill. ≥ 8144 lx (S2 Table). Conversely, the lowest CATE estimate of 0.036 (95% CI: -0.12–0.20) was found under Hum. < 79% and Tem. ≥ 24 °C (S1 Table). These findings suggest that environmental factors, such as humidity, light, and temperature, influence the effectiveness of NaCl treatment on °Brix. Additionally, we examined the relationship between cultivation periods and the subgroups derived from the Causal Tree. Cultivation period D was classified into only one subgroup due to its small sample size, whereas datasets from other growing periods were distributed across multiple subgroups (Fig 4). By contrast, each Causal Tree subgroup included samples from multiple cultivation periods (Table 4). This indicates that the subgrouping based on the Causal Tree was not merely a result of seasonal classification but rather driven by environmental and contextual differences.

**Fig 4.**
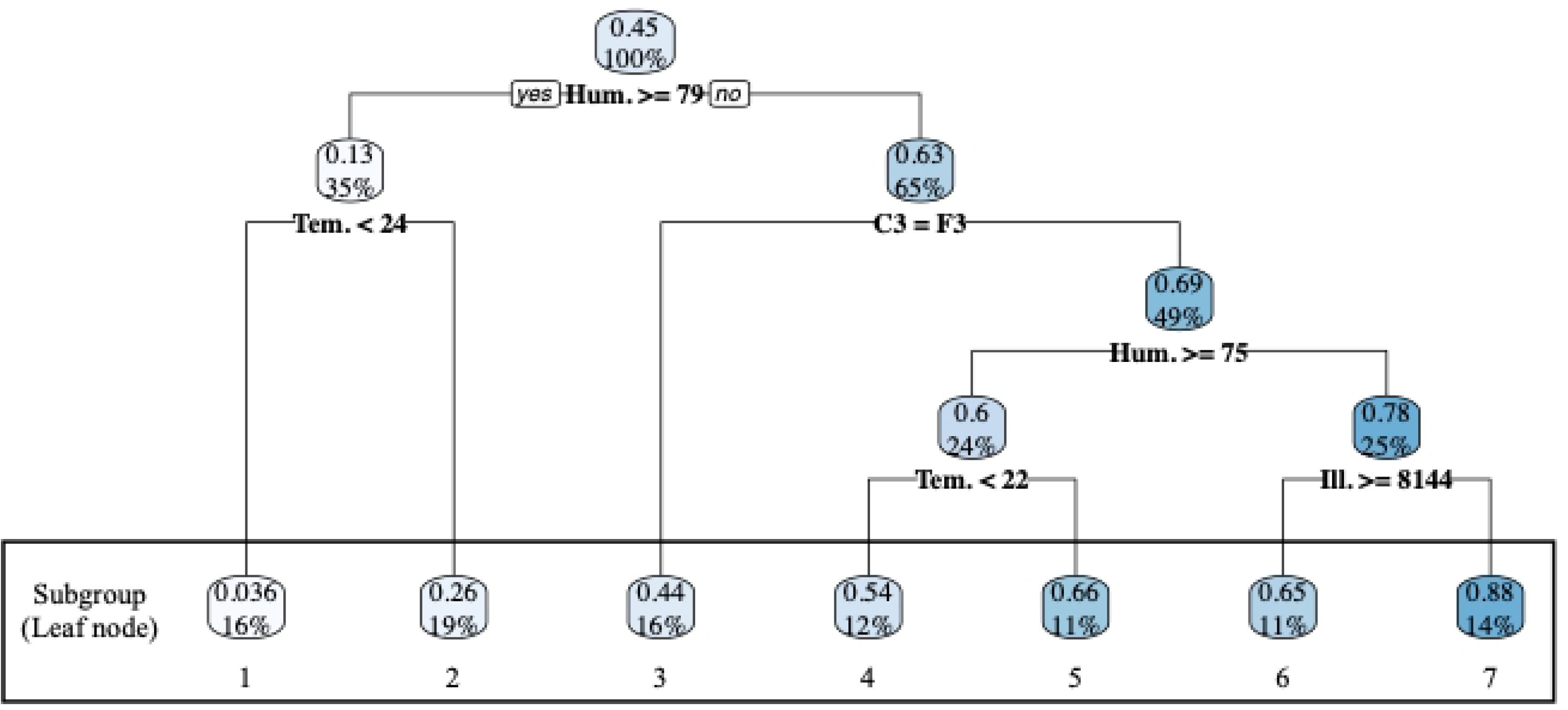
Causal Tree illustrating factors influencing tomato °Brix. Each node represents a splitting variable, with data partitioned based on the threshold values. The subgroups indicate the classified groups and their respective proportions. The values above and below each node denote the CATE and the proportion of Dataset B, respectively.

**Supplementary table 2.** Subgroup-Specific CATE derived from Causal Tree splits based on environmental conditions.

| Subgroup | n (W=1 / W=0) | $\hat{y}(T=1)$ | $\hat{y}(T=0)$ | CATE | 95% CI |
| --- | --- | --- | --- | --- | --- |
| 1 | 45 / 47 | 0.8444 | 0.8085 | 0.0359 | [-0.12, 0.20] |
| 2 | 61 / 51 | 0.3770 | 0.1176 | 0.2594 | [0.10, 0.41] |
| 3 | 43 / 50 | 0.7209 | 0.2800 | 0.4409 | [0.25, 0.63] |
| 4 | 34 / 37 | 0.6765 | 0.1351 | 0.5413 | [0.34, 0.74] |
| 5 | 33 / 32 | 0.7273 | 0.0625 | 0.6648 | [0.48, 0.85] |
| 6 | 33 / 30 | 0.8485 | 0.2000 | 0.6485 | [0.45, 0.85] |
| 7 | 40 / 42 | 0.9750 | 0.0952 | 0.8798 | [0.77, 0.99] |

Each row represents a subgroup defined by a combination of covariate conditions identified by the Causal Tree. Treatment ( $W = 1$ ) and control ( $W = 0$ ) groups were compared within each subgroup. CATE represents the difference in outcome means between treated and control groups, with 95% confidence intervals (CI) calculated using standard error estimates.

## Discussion

We examined the effects of a NaCl treatment on fruit quality across different cultivation periods. First, we confirmed that the NaCl treatment affected fruit quality throughout all cultivation periods. Second, the effects on fruit quality differed depending on the cultivation period. Third, we evaluated the conditional effects of treatments by incorporating environmental factors as covariates and found that humidity had the most significant impact on the treatment effects.

NaCl treatment increases the osmotic potential of the nutrient solution, thereby suppressing water uptake by plants and reducing fruit water content. This consequently leads to a relative increase in the concentration of soluble sugars within the fruit. Ladewig et al. [14] reported a significant increase in sugar content (°Brix) in response to elevated NaCl concentrations. Among environmental factors, humidity is a key determinant of plant water stress. High humidity conditions alleviate the reduction in water potential and yield under salt stress, likely by suppressing excessive transpiration and maintaining leaf turgor [20,28]. Furthermore, Romero-Aranda et al. [28] reported that mist treatment under saline irrigation improved leaf water potential, at least doubled tomato yield, and reduced Na⁺ accumulation in leaves. In addition, Yu et al. [23] demonstrated that low vapor pressure deficit during the fruit expansion stage increased fruit water content, potentially reducing °Brix. In our study, humidity was the most influential factor in determining the effect of mild NaCl treatment (EC4, 25 mM) on fruit sugar content. The relative humidity during the cultivation periods ranged from 60% to 85%, and a threshold below 79% was associated with significant sugar content enhancement (Fig 4, Table 2). These findings are consistent with those of previous reports, indicating that the physiological effects of water stress are more pronounced under low-humidity conditions.

Causal Tree analysis was used to explore the factors influencing the variation in the effect of NaCl treatment on tomato fruit sugar content (Fig 4). The analysis revealed that relative humidity was the most influential factor, followed by the truss position, indicating that both environmental conditions and plant physiological status significantly affected the magnitude of the treatment effect. Specifically, the most effective conditions for increasing °Brix were identified as fruits from the third truss grown under an average relative humidity of ≤ 75% and a cumulative light intensity ≥ 8,144 lx. These conditions corresponded to subgroup 7, which included fruits from cultivation periods B, G, and H, suggesting that the effect was not due to a specific cultivation period but to a reproducible environmental combination.

However, differences in °Brix and the effects of the NaCl treatment were observed even among fruits cultivated under the same conditions, which may be attributed to variations in the sink– source ratio. In this experiment, the number of fruit sets was not limited, which likely caused variability in the fruit load per plant and consequently affected sugar accumulation. Previous reports support this explanation. Prudent et al. [29] reported co-localized quantitative trait loci with opposing effects on fruit weight and sugar content, suggesting a genetic trade-off under high sink demand. Jan et al. [30] showed that increasing the leaf-to-fruit ratio increased sugar and sucrose concentrations in the phloem sap. These findings indicate that the balance between assimilate supply and sink demand is central to sugar accumulation. Future analyses should incorporate these parameters to investigate further how both environmental and plant-level factors regulate sugar content and yield.

This study had several limitations. First, data were collected using a single tomato cultivar from a single greenhouse site. Although the dataset was highly accurate, the findings may not be generalizable to other cultivars, greenhouse facilities, or environmental settings. Future studies should validate these results using datasets from multiple sites and cultivars under various climatic and soil conditions. Second, while the Causal Tree method offers high interpretability and facilitates the exploration of treatment effect heterogeneity, its results may be sensitive to hyperparameter selection and the limited sample size within subgroups. Third, this study was conducted as an exploratory pilot study, and the sample size used in the analysis was relatively small. Therefore, the specific patterns and associations between variables observed in the results of the Causal Tree analysis may be specific to this dataset, and their generalizability and robustness need to be verified in further large-scale studies. In particular, the stability of the Causal Tree using Honest Estimation and a detailed robustness evaluation using the bootstrap method are issues to be addressed in the future. Finally, the outcome variables were classified using a provisional Brix threshold of 6% based on the distribution of the observed data. Depending on specific research objectives or practical quality standards in future studies, this threshold may require adjustment.

Furthermore, while this study focused on °Brix as an indicator of fruit quality, future research should also address the potential trade-offs between sugar enhancement and yield stability, as well as the risk of salt toxicity under varying salinity levels and across different tomato varieties.

## Conclusions

In this study, tomato plants were cultivated year-round using hydroponics across nine different cultivation periods (A–I), each with a distinct sowing date. We applied a NaCl treatment (25 mM) to examine how salinity stress influenced fruit sugar content (°Brix) across the cultivation periods. The sugar content ranged from 4.8 to 6.2 °Brix in the control groups and from 5.4 to 7.6 °Brix in the NaCl-treated groups, with consistently higher values in the NaCl-treated plants.

The effect of NaCl treatment on the sugar content varied significantly depending on the cultivation period. The greatest effect was observed during cultivation period H, which differed from periods F (the highest sugar content in controls) and E (the lowest sugar content). This suggests that the optimal conditions for enhancing fruit sugar content in untreated plants do not necessarily align with those that maximize the effect of NaCl treatment, implying that unique environmental factors influence treatment efficacy. A Causal Tree analysis indicated that relative humidity was the most influential factor, followed by the fruit truss stage and cumulative illuminance.

Our findings demonstrate that the Causal Tree approach can effectively identify the environmental conditions necessary to maximize the efficacy of NaCl treatment for increasing fruit sugar content, thereby guiding precise environmental control strategies for tomato cultivation in controlled environments such as greenhouses or plant factories.

By enabling the estimation of heterogeneous treatment effects, this causal inference method allows us to go beyond simple correlations and identify conditions under which NaCl treatment has a causal effect on sugar accumulation. This insight is essential for optimizing cultivation in diverse and variable environments.

## Acknowledgments

We sincerely thank Dr. Yasunaga Iwasaki of Meiji University for his valuable advice on salt-stress treatment. We also thank Ms. Chisato Goto for her helpful support throughout this study.

